# Liquid Lung Rest During Extracorporeal Life Support in a Porcine Model of Acute Lung Injury

**DOI:** 10.1101/2025.10.17.683108

**Authors:** David G. Blauvelt, Anne Hesek, Michael Golecki, Prasad Nithianandam, Gianna M. Parrish, James Keith, Lynell Jones, Kelly Massa, Scott L. Weiss, Thomas Shafffer

## Abstract

**Rationale:** In patients with severe lung injury, extracorporeal life support can enable lung rest, but the optimal strategy is unknown.

**Objective:** To compare liquid lung rest versus standard gas rest ventilation in a porcine model of acute lung injury and extracorporeal life support.

**Methods:** Twenty neonatal pigs with lung injury after systemic injection of oleic acid were placed on extracorporeal life support. Pigs were ventilated with standard rest settings (peak inspiratory pressure: 20 cmH_2_O; end expiratory pressure: 10 cmH_2_O; rate: 10 breaths/minute) and randomized to receive no additional therapy (gas ventilation) or 5 mL/kg perfluorooctylbromide instilled endotracheally (liquid lung rest). The study continued for 4 hours with hourly pulmonary compliance measurements. After euthanasia, lung samples were taken for histologic analysis. Inflammation was quantified via blood and lung tissue cytokines.

**Measurements and Main Results:** All pigs achieved significant lung injury; static compliance decreased by a mean of 54%. After 4 hours of lung rest, static compliance was higher with liquid lung rest compared to gas ventilation (0.88±0.05 mL/kg/cmH_2_O versus 0.57±0.05, p<0.001).

Histology revealed 2.1-fold greater airspace in dependent lung regions with liquid lung rest (p<0.001) and a reduced airspace heterogeneity index (LLR: 0.071±0.004, gas: 0.095±0.010, p=0.04), suggesting more uniform alveolar recruitment. Cytokine analysis demonstrated a 3.7-fold decrease in tissue interleukin-10 levels with liquid lung rest (p=0.02).

**Conclusions:** Liquid lung rest improved pulmonary compliance, achieved better alveolar recruitment, and decreased tissue interleukin-10 levels compared to standard gas ventilation. This strategy may enhance lung recovery in acute lung injury requiring extracorporeal life support.

## 1. Introduction

Severe acute respiratory distress syndrome (ARDS) has a mortality as high as 46% (1). It results from acute lung injury causing fluid, protein, and inflammatory cell leakage into alveoli, leading to hypoxemia, acidosis, and, in severe cases, multiorgan failure and death. Extracorporeal life support (ECLS) is an advanced therapy for severe ARDS that may improve outcomes (2-5). ECLS circulates blood through an oxygenator for gas exchange, allowing lower mechanical ventilation pressures, potentially aiding lung recovery. However, ECLS carries substantial risks (6), underscoring the need for concurrent therapeutic strategies that promote lung healing and shorten ECLS duration.

Ventilation strategies aiming for uniform alveolar opening may limit ongoing lung damage, but ARDS involves heterogenous injury – alveolar collapse in some regions and overdistension in others (7). Thus, lung rest using conventional gas ventilation has produced inconsistent results, reflective of the variability of the disease (8-14). A method enabling homogeneous alveolar opening to their resting state may accelerate lung recovery on ECLS.

Liquid lung rest (LLR) is a strategy that may offer benefits over conventional gas ventilation. Liquids can more evenly recruit lung tissue at lower pressures, reach dependent alveoli prone to collapse, and enhance pulmonary clearance by displacing bacteria, mucus, and inflammatory cells to the surface (15, 16). Liquids can also enable targeted drug delivery to the lungs, including antibiotics, mucolytics, or anti-inflammatory agents (17). Despite these potential benefits, LLR has not been studied as a therapeutic strategy for ARDS requiring ECLS. Notably, because ECLS performs oxygenation and ventilation, LLR can focus on minimizing ventilator-induced lung injury – including barotrauma, atelectrauma, and biotrauma – and promote lung healing, rather than facilitating gas exchange. The aim of this study was to compare LLR versus lung rest with conventional gas ventilation using a porcine model of acute lung injury and ECLS, assessing pulmonary compliance, alveolar recruitment, and inflammation.

## 2. Methods

The overall experimental design is illustrated in Figure 1. All methods were approved by the Institutional Animal Care and Use Committee (IACUC) at Nemours Children’s Health.

**Figure 1:**
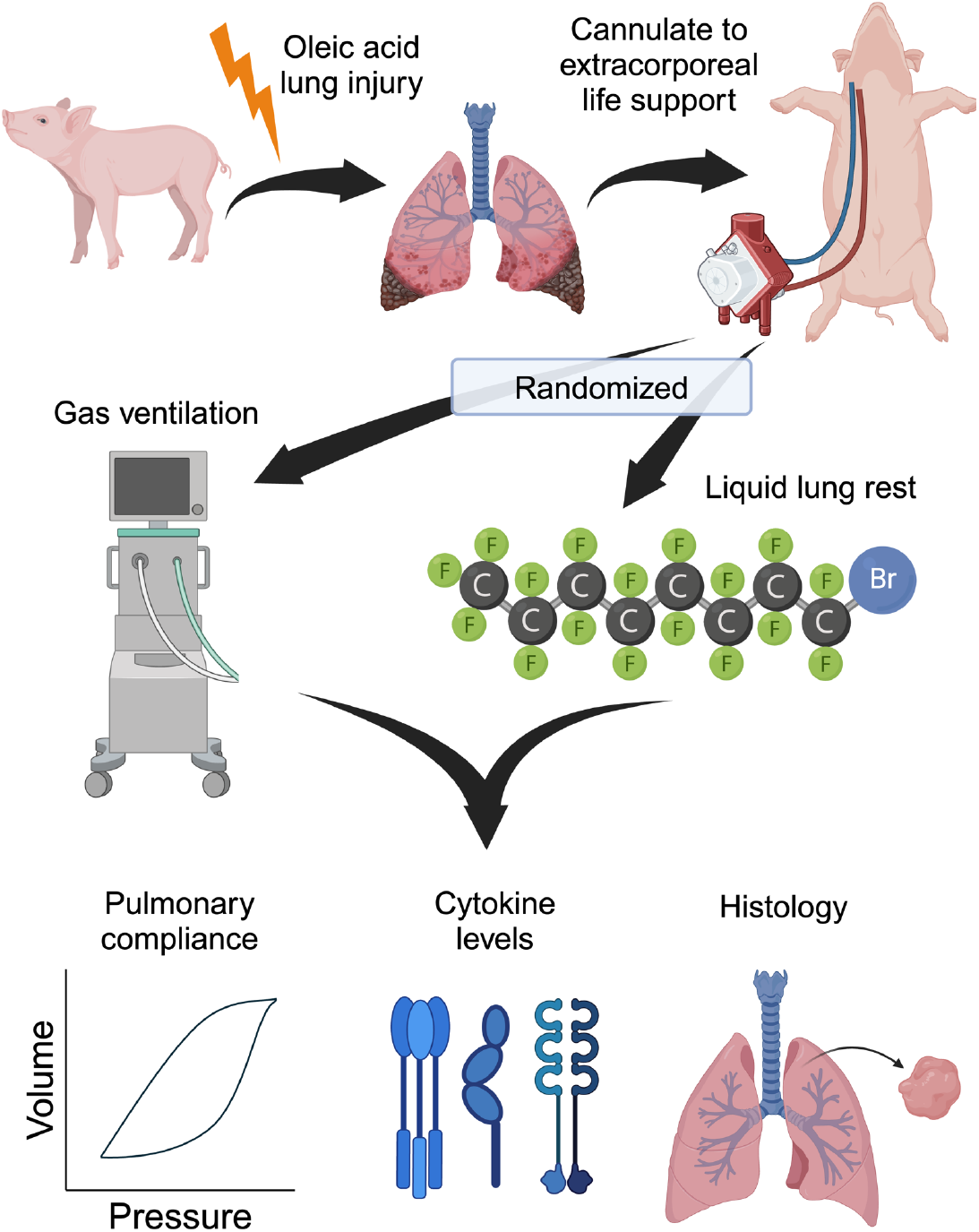
Experimental design. Neonatal pigs underwent lung injury with oleic acid and were cannulated to veno-arterial extracorporeal life support (VA-ECLS). They were then randomized to receive either gas ventilation with rest settings or liquid lung rest. In the liquid lung rest group, 5 mL/kg of perfluorooctylbromide (PFOB) was instilled into the lungs. The outcomes measured were pulmonary compliance, cytokine levels, and histologic analysis of alveolar recruitment.

### Animal Preparation

Neonatal pigs (Age: ∼10 days old; Weight ∼4 kg) underwent anesthetic induction with intramuscular ketamine-acepromazine-xylazine (KAX). A central venous catheter (5.5 French, triple lumen, Teleflex Medical) and arterial catheter (5 French, single lumen, Vygon) were placed into the left jugular vein and left carotid artery, respectively. Anesthesia was maintained with continuous infusions of propofol and ketamine. The pigs were intubated with a 3.5 cuffed endotracheal tube and placed on mechanical ventilation (Servo-i, Getinge): volume control (VC); tidal volume: 8 mL/kg; positive end expiratory pressure (PEEP): 5 cmH_2_O; fraction of inspired oxygen (F_i_O_2_) = 0.50. Throughout the experiment the pigs had continuous monitoring of their heart rate, respiratory rate, blood pressure, oxygen saturation, and temperature.

### Lung Injury Protocol

An oleic acid injury protocol was used to model indirect acute lung injury, as previously demonstrated (18-21). Oleic acid aliquots (0.02 mL/kg, #AA3199706, Fisher Scientific) were mixed with an equal quantity of sterile 0.9% saline. The oleic acid was emulsified by vortexing the mixture, diluted with an additional 2 mL of freshly drawn blood, and injected into the central venous catheter. A total of four aliquots (0.08 mL/kg total oleic acid dose) were administered, allowing for 5-10 minutes of recovery between each administration. Transient hypotensive episodes occasionally occurred and were treated with intermittent boluses of norepinephrine (1 mcg/kg). Prior to injury and after the 4^th^ aliquot, blood gases were drawn to quantify the ratio of arterial partial pressure of oxygen (P_a_O_2_) to F_i_O_2_ (P/F ratio).

Additionally, baseline and post-injury static pulmonary compliances (C_stat_) were measured. Sufficient injury was defined as a 50% decrease in C_stat_ or a P/F ratio of less than 200. If these criteria were not met, up to two additional oleic acid doses of 0.02 mL/kg were given (maximum 0.12 mL/kg total dose).

### Extracorporeal Life Support Initiation

Following the oleic acid lung injury, the pigs were cannulated to ECLS. An arterial cannula (6-8 French, Biomedicus, Medtronic) was placed in the right carotid artery and advanced to the aortic arch. A venous cannula (8-10, French, Biomedicus, Medtronic) was placed in the external jugular vein and advanced to the right atrium. The ECLS circuit consisting of a roller pump (S3, Sorin), oxygenator (D101, LivaNova), and water heater (SMS-5000, Seabrook Medical Systems) was primed with freshly drawn citrated donor pig blood, calcium chloride (300 mg), unfractionated heparin (600 IU), and sodium bicarbonate (10 mEq). Once the circuit was connected to the cannulas, blood flow was increased to a target of 100 mL/kg/min. Oxygen sweep gas was initiated at 0.4 mL/min and titrated to an arterial pH of 7.35-7.45 and arterial CO_2_ of 35-45 mmHg. Unfractionated heparin was used to maintain anticoagulation: 150 IU/kg bolus immediately prior to cannulation and an infusion titrated to an activated clotting time of 200-250 s (Hemochron Signature Elite, Werfen).

### Experimental Protocol

The experimental protocol began once the pigs were established on stable ECLS settings. Pigs were randomized to one of two treatments – LLR or gas ventilation – by drawing group assignments from a box containing an equal number of slips labeled for each study arm. In both groups, pigs were supported with standard lung rest settings: pressure control ventilation (PCV); positive inspiratory pressure (PIP): 20 cmH_2_O; PEEP: 10 cmH_2_O; respiratory rate: 10 breaths/min; F_i_O_2_: 0.40. In the LLR group, pigs also received 5 mL/kg of liquid perfluoroctylbromide (PFOB), instilled directly into the endotracheal tube. The pigs were rotated side to side and 45° reverse Trendelenburg to allow for uniform dosing throughout the lung fields. The experimental protocol continued for 4 hours. Each hour, pulmonary compliance was measured, and blood samples were drawn for titration of the ECLS circuit, anticoagulation management, and measurement of cytokine levels. 1 mL/kg of PFOB was added hourly to account for evaporative losses.

### Tissue Harvest and Histological Preparation

At the end of the study, pigs were humanely euthanized with Euthasol (phenytoin/pentobarbital). With the trachea clamped at the expiratory volume, the chest cavity was exposed via thoracotomy. The main pulmonary artery was cannulated, and the lungs were perfused with refrigerated phosphate-buffered saline (PBS) until the effluent cleared, taking care to minimize the perfusion pressure to <30 cmH_2_O. The right lung was used for histologic analysis, and the left lung was used for tissue cytokine measurements. Samples (approximately 1 cm^3^) were taken from the anterior and posterior portions of each lobe (right lung: upper/middle/lower lobes; left lung: upper/lower lobes). Samples for tissue cytokine quantification were snap frozen in liquid nitrogen. Samples for histology were preserved in 10% formalin, embedded in paraffin, sectioned into 5 µm sections, mounted on slides, and stained with hematoxylin and eosin (H&E).

### Image Analysis

Stained tissue samples were imaged using widefield light microscopy (EVOS M7000, Invitrogen) and analyzed using custom Matlab scripts. Regions of interest were manually identified to include the alveolar parenchyma for analysis while excluding large airways and vasculature. The regions of interest were then converted to binary images such that each pixel was classified as either air or tissue. The airspace density was calculated as the fraction of pixels that were identified as air. Additionally, a heterogeneity index describing the interregional variability of alveolar expansion was calculated. The heterogeneity index was defined as the standard deviation of airspace density across 24 samples within a single animal.

### Plasma and Tissue Cytokine Levels

Blood samples were drawn into lithium heparin Vacutainers (Becton Dickinson) before and after lung injury as well as hourly during the experimental protocol. Blood was centrifuged at 2000 g for 10 minutes to separate the plasma from the cellular components and frozen on dry ice. Plasma cytokine levels were measured using the Luminex multiplex assay kit for porcine cytokines (Porcine ProcartaPlex Simplex Kits, Thermo Fisher). For tissue cytokine levels, 50-100 µg samples of frozen lung tissue were homogenized in cell lysis buffer (Fisher Scientific) using an Omni Bead Mill Homogenizer (6 m/s for 2 x 15 s cycles). The homogenized tissue in lysis buffer was centrifuged (16,000 g x 10 minutes), and the supernatant was used for cytokine measurement using the Luminex multiplex assay. Tissue cytokine levels were normalized to the total protein content of the supernatant, which was measured with the Bicinchoninic Assay (BCA).

### Statistical Analysis

Statistical analysis was performed using GraphPad Prism (version 10). Hourly measurements (compliance and plasma cytokine levels) were compared using a repeated measures analysis of variance (ANOVA), incorporating factors of treatment (gas versus LLR) and repeated measures on time. Bonferroni’s multiple comparisons test was used to compare the treatment groups at each hour and to compare hourly measurements to the baseline values within each treatment group. Measurements collected from various lung zones (airspace density, tissue cytokines) were also compared with ANOVA, incorporating repeated measures on lung zone. Bonferroni’s test compared airspace density between treatment groups at each lung zone. Heterogeneity index was compared using a 2-tailed unpaired t-test. For both plasma and tissue cytokine levels, the measured values were log10 transformed prior to statistical analysis. To determine sample size, an interim analysis was performed after 12 pigs (6 per group) using C_stat_ data. The mean ± standard deviation of C_stat_ at 4 hours was 0.83±0.17 mL/kg/cmH_2_O and 0.58±0.20 mL/kg/cmH_2_O in the LLR and gas groups, respectively. Thus, 10 pigs per group were needed to achieve 81% power to detect a significant difference in C_stat_ between groups at 4 hours at an α=0.05. Descriptive statistics are presented as mean ± standard error of the mean unless otherwise specified. P-values <0.05 were deemed to be statistically significant.

## 3. Results

Twenty pigs (10 per treatment arm) completed the study protocol and were included in the final analysis. Four pigs that died during the lung injury protocol or were unable to be cannulated to ECLS (i.e. prior to treatment randomization) were excluded. Baseline and clinical characteristics are presented in Table 1. There were no significant differences in baseline characteristics between the gas and LLR groups. Across all pigs, both C_stat_ and dynamic pulmonary compliance (C_dyn_) had a significant decrease from baseline after the oleic acid injury (C_stat_: 1.24±0.03 mL/kg/cmH_2_O to 0.57±0.02 mL/kg/cmH_2_O, p<0.001 [Paired 2-tailed t-test]; C_dyn_: 0.95±0.02 mL/kg/cmH_2_O to 0.490±0.01 mL/kg/cmH_2_O, p<0.001). Lung injury trended toward more severe (lower C_stat_ after injury) in the LLR group, but this difference was not statistically significant. P/F ratio was 182±12 across all pigs, with no difference between the two groups.

**Table 1:** Baseline and clinical characteristics of the subjects. Data are presented as mean ± standard deviation Cdyn = dynamic compliance; Cstat = static compliance; PaCO2 = partial pressure of carbon dioxide; P/F ratio = ratio of partial pressure of arterial oxygen to fraction of inspired oxygen

|  | Gas Ventilation | Liquid Lung Rest | P-value |
| --- | --- | --- | --- |
| Age (days) | 9.8 ± 1.3 | 10.2 ± 1.2 | 0.49 |
| Weight (kg) | 4.3 ± 0.4 | 4.2 ± 0.4 | 0.65 |
| C<sub>dyn</sub>(mL/kg/cmH<sub>2</sub>O) |  |  |  |
| Pre-injury | 0.97 ± 0.12 | 0.93 ± 0.08 | 0.40 |
| Post-injury | 0.51 ± 0.06 | 0.47 ± 0.03 | 0.17 |
| C<sub>stat</sub>(mL/kg/cmH<sub>2</sub>O) |  |  |  |
| Pre-injury | 1.25 ± 0.14 | 1.22 ± 0.14 | 0.77 |
| Post-injury | 0.60 ± 0.09 | 0.54 ± 0.03 | 0.06 |
| Oleic Acid Dose (mL/kg) | 0.09 ± 0.02 | 0.09 ± 0.01 | 0.29 |
| Post-Injury Blood Gas |  |  |  |
| pH | 7.25 ± 0.07 | 7.25 ± 0.08 | 0.93 |
| PaCO<sub>2</sub> (mmHg) | 53.2 ± 15.9 | 52.7 ± 8.5 | 0.93 |
| P/F ratio | 183 ± 60 | 181 ± 47 | 0.93 |
| ECMO Blood Flow Rate (mL/kg/min) | 97.3 ± 8.4 | 97.4 ± 5.7 | 0.99 |

In the gas group, neither C_stat_ nor C_dyn_ significantly changed over the course of the 4-hour experiment, relative to the post-injury compliance (Figure 2). By contrast, in the LLR group, both C_stat_ and C_dyn_ increased relative to post-injury at all time points. At the end of the study (hour 4), C_stat_ in the LLR group had increased to 0.88±0.05 mL/kg/cmH_2_O (versus LLR post-injury: 0.54±0.01 mL/kg/cmH_2_O, p=0.003; versus gas at hour 4: 0.57±0.05 mL/kg/cmH_2_O, p<0.001), and C_dyn_ had increased to 0.62±0.03 mL/kg/cmH_2_O (versus LLR post-injury: 0.47±0.01 mL/kg/cmH_2_O, p=0.004; versus gas at hour 4: 0.49±0.04 mL/kg/cmH_2_O, p=0.014).

**Figure 2:**
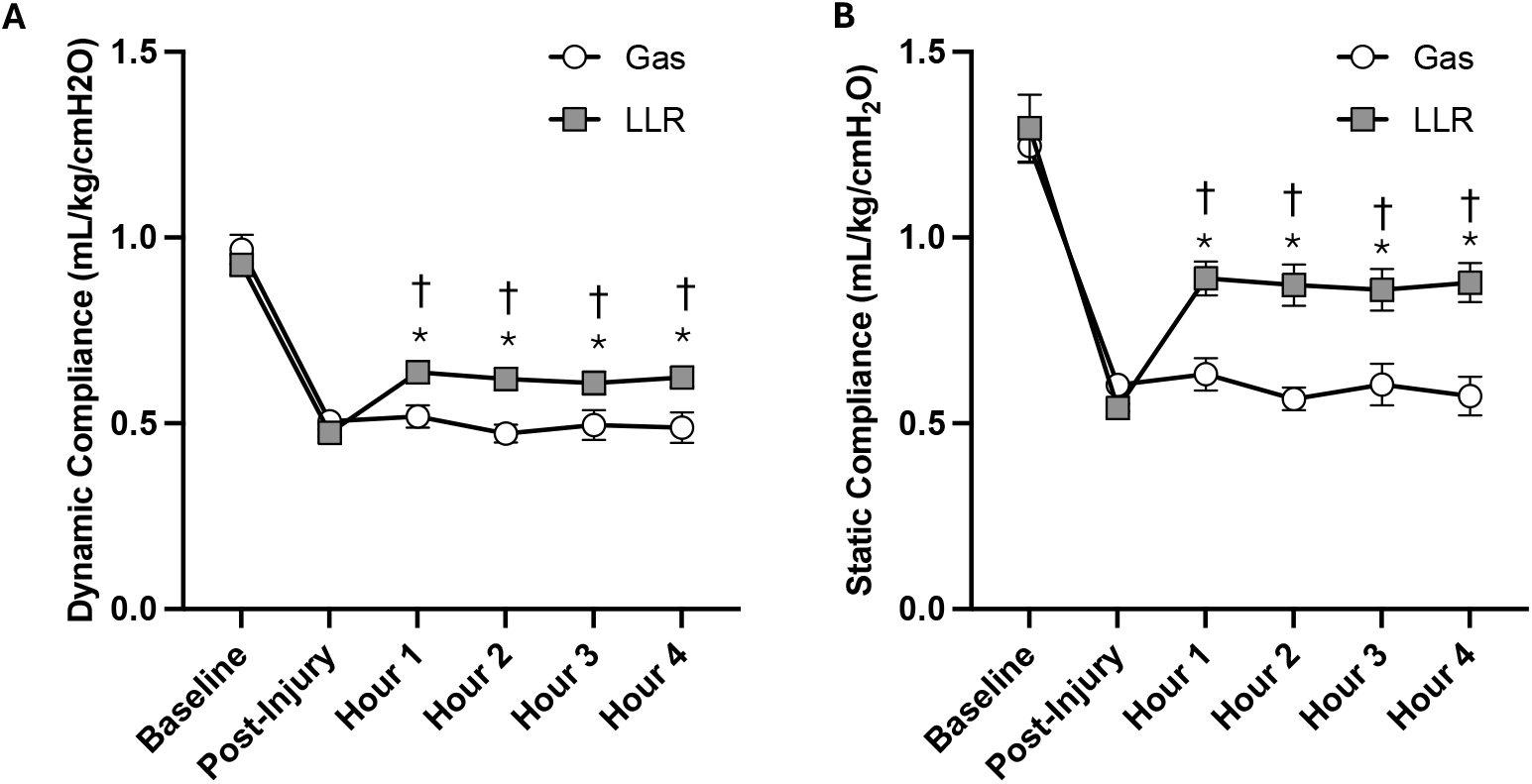
Pulmonary compliance data. (A) Dynamic and (B) static compliance at baseline, immediately following the acute lung injury, and hourly during the experimental protocol. Markers denote mean compliance. Error bars represent standard error. p<0.05, ⁎: versus post-injury value; †: versus gas ventilation

Following the end of the experimental protocol, a necropsy was performed, and samples were taken from the right lung for histological analysis (Figure 3). Gross inspection of the lung revealed significant lung injury, disproportionately affecting the dependent portions of the lung. Microscopic examination showed that the alveoli were subjectively more open in the LLR group versus the gas group, especially in the dependent areas. We next quantified the airspace density in the lung tissue (Figure 4). Samples were categorized as originating from one of six zones, with zone 1 representing the most non-dependent region, and zone 6 denoting the most dependent area (Figure 4A). Overall mean airspace was not different between groups (LLR: 0.29±0.01, gas: 0.27±0.01, p=0.43; Figure 4B), but there was a highly significant interaction between lung sample location and treatment (F(11,198) = 4.95, p<0.001). Post-hoc comparison of the airspace density within each lung zone (Figure 4C) demonstrated a significantly airspace density only in zone 6 (LLR: 0.34±0.02, gas: 0.16±0.03, p<0.001), suggesting that the effect of LLR on alveolar opening was primarily on the most dependent area of the lung. The heterogeneity of airspace density throughout the lung (heterogeneity index) was then quantified to determine whether LLR influenced the uniformity of alveolar opening (Figure 4D). The LLR group had a lower heterogeneity index (LLR: 0.071±0.004, gas: 0.095±0.010, p=0.04) suggesting more uniform alveolar recruitment.

**Figure 3.**
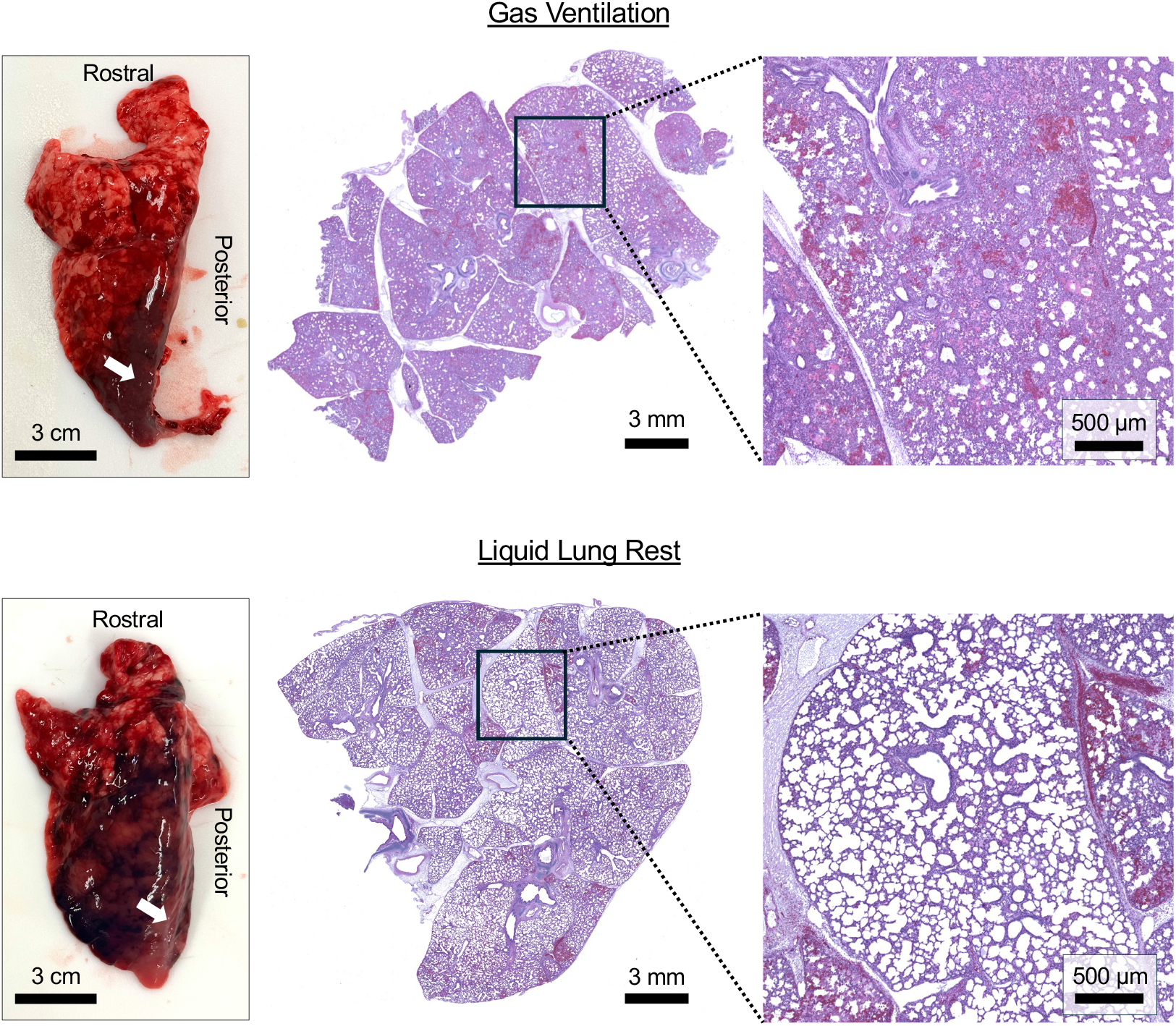
Lung histology. Left panels: Gross histologic examination demonstrating severe lung injury in both the gas ventilation (top) and liquid lung rest (bottom) groups. Middle and right panels: Representative microscopic images of lung samples taken from a similar location in the dependent portion of the lung (white arrows on gross histology panels). Subjectively, the liquid lung rest group had increased and more homogenous alveolar recruitment.

**Figure 4.**
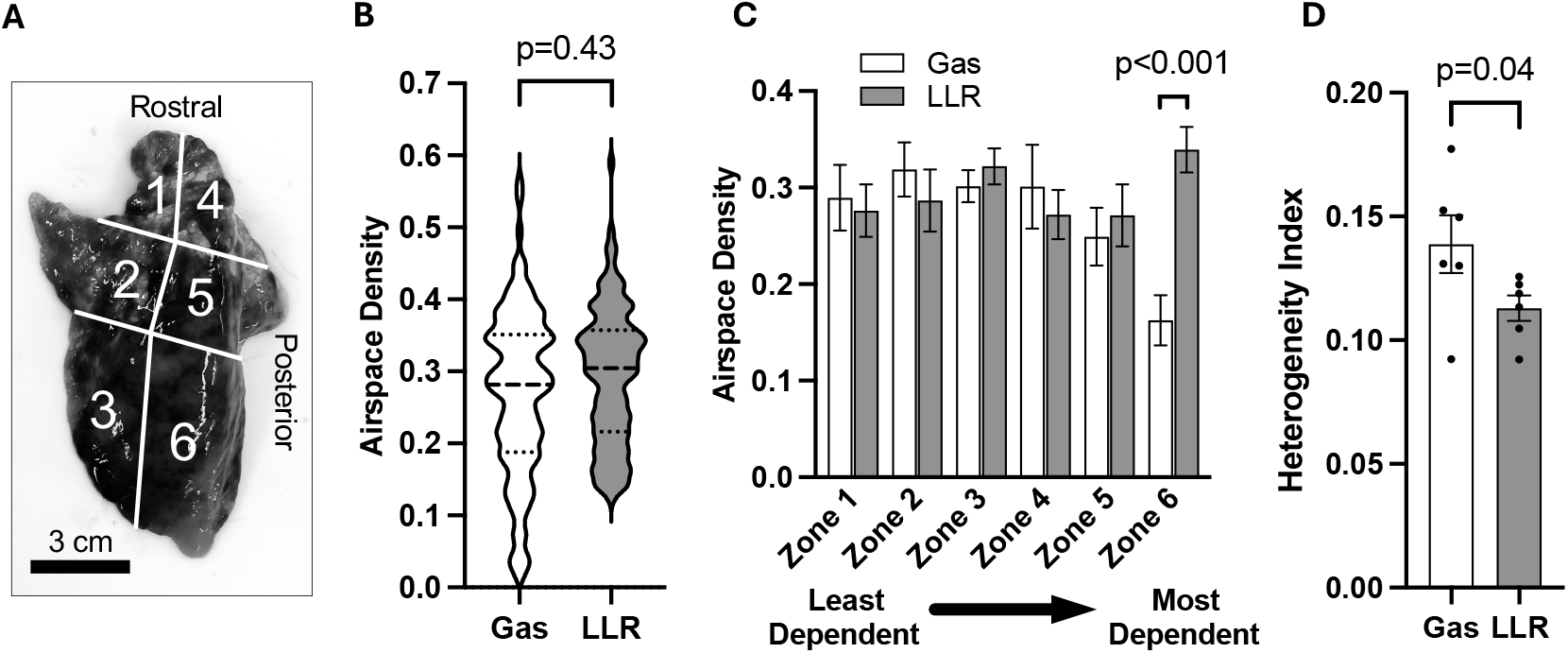
Quantitative analysis of alveolar recruitment. (A) The lung samples were taken from one of size lung regions. Zone 1 was the anterior part of the upper lobe (most non-dependent region). Zone 6 is the posterior part of the lower lobe (most dependent region). (B) Violin plot showing the distribution of airspace density values measured across all pigs and samples (n = 10 pigs per group, 24 samples per pig). Width represents the relative frequency distribution. Dashed lines denote the 25^th^, 50^th^, and 75^th^ percentiles. (C) Airspace density when evaluated by individual zone. Bar graph shows mean airspace density. Error bars represent standard error. (D) Heterogeneity index denoting the uniformity of airspace density across the 24 samples within the same pig. Points represent values from individual pigs. Bar graph shows mean heterogeneity index. Error bars represent standard error.

Plasma samples were taken at baseline, after injury, and hourly on ECLS to measure cytokine levels throughout the experiment (Figure 5). Six cytokines were evaluated: interleukin-1beta (IL-1β), IL-4, IL-6, IL-8, IL-10, and tumor necrosis factor alpha (TNF-α). One pig in the gas ventilation group was excluded from the primary cytokine analyses due to outlier results, with plasma cytokine levels that were 2-3 orders of magnitude higher than all the other pigs. Among the remaining 19 pigs, there were no significant differences in plasma cytokine levels between groups at any time point. A sensitivity analysis (Figure S1, Supplemental Material) including the outlier pig found the same results, with the exception that TNF-α was lower in the LLR group at hour 3 (LLR: 408±134 pg/mL, gas: 1492±923 pg/mL, p=0.05).

**Figure 5.**
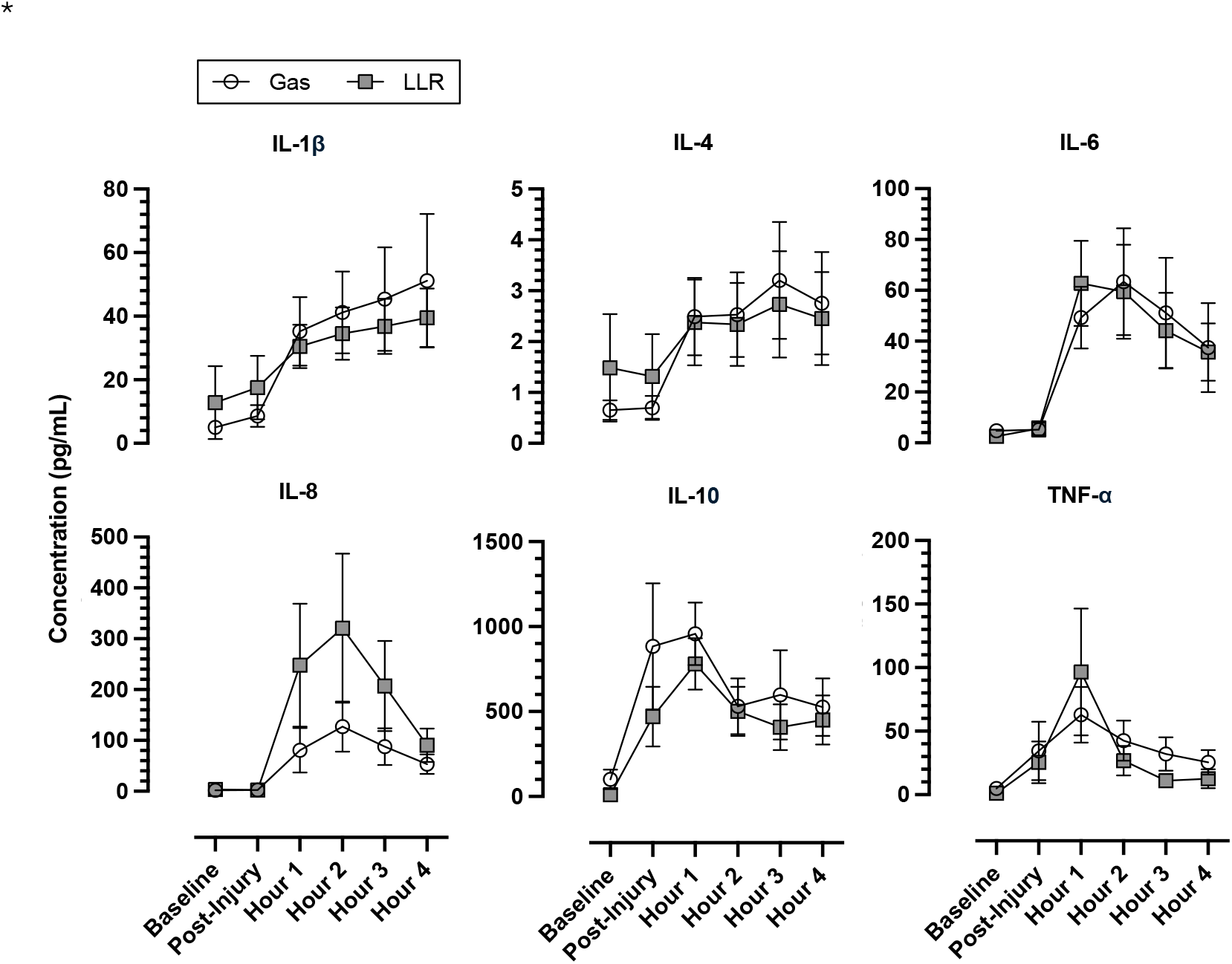
Plasma cytokine levels. Plasma samples were drawn at baseline, after lung injury, and hourly while on ECLS. One pig in the gas ventilation group had extremely high cytokine levels and was excluded as an outlier. There were no statistical differences between the groups at any timepoint. Markers denote mean cytokine levels. Error bars denote standard error.

Cytokine levels in lung tissue homogenates were also measured. To account for different sizes of tissue, cytokine levels were normalized to the total protein amount and expressed as a fraction of total protein (Figure 6). All cytokines had levels that trended lower in the LLR group, but differences were only significant for IL-10. The fraction of IL-10 was 3.7 times lower in the LLR group (LLR: 6.0 10^-9^±1.2 10^-9^, gas: 2.2 10^-8^±1.0 10^-8^, p=0.02).

**Figure 6.**
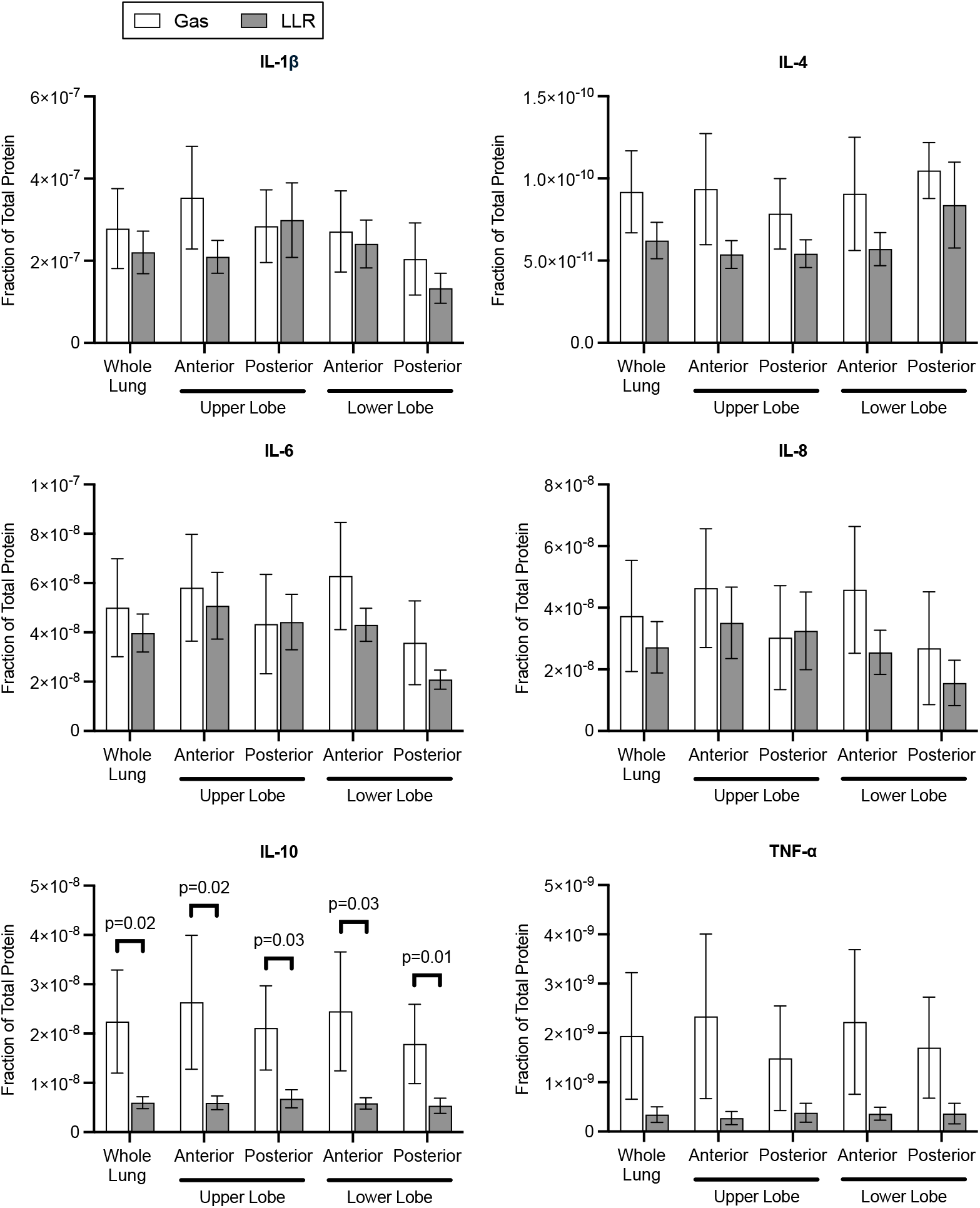
Lung tissue cytokine levels. Lung tissue samples were taken at the end of the study and homogenized to quantify cytokine levels. There was a statistically significant decrease in tissue IL-10 level in the LLR group. There were no statistically significant differences in other cytokine levels. Bar graphs show mean cytokine levels expressed as a fraction of total protein. Error bars denote standard error.

## 4. Discussion

The ability to reduce mechanical ventilation to allow for lung rest and recovery is a key benefit of ECLS therapy. In this study, we evaluated LLR as an adjunct to conventional lung rest using low ventilator pressures. Pulmonary compliance improved significantly in the LLR group compared to gas ventilation. Histology also showed improved alveolar opening in dependent lung regions, which are disproportionately collapsed in ARDS. This corresponded with reduced cytokine levels, most notably lung tissue IL-10 levels.

Recent studies have sought to identify optimal ventilation strategies for lung rest during ECLS, examining tidal volume, PEEP, driving pressure, and F_i_O_2_. However, the findings of a multitude of retrospective and prospective clinical studies have been contradictory. For instance, although three studies showed that high F_i_O_2_ is associated with increased mortality (9, 22, 23), a later meta-analysis found no significant association of F_i_O_2_ and mortality (24). Similarly, while some studies have found better outcomes with higher PEEP (8, 25), others reported worse outcomes (9, 26) and several found no difference (12, 24). Thus, the optimal ventilation strategy for patients on ECLS remains controversial. Perhaps large, randomized trials will shed light on an optimal strategy, but given the variability in findings, any statistically significant effects are likely to be small. Our approach of using liquid to recruit collapsed alveoli in a more uniform manner introduces a new therapeutic strategy that may yield more consistent benefits.

Although LLR for ECLS is a novel strategy, the related concept of using perfluorocarbons such as PFOB for liquid ventilation has been studied for decades (27-29). In both human and animal studies, liquid ventilation has been shown to improve compliance and gas exchange (18, 30-32). However, a 2006 multicenter randomized trial failed to show improved outcomes in adult ARDS (33). Although liquid ventilation fell out of favor following that study, more recent efforts have examined the possible benefits of liquid ventilation in certain contexts, including younger patients and those with severe disease (29, 33, 34). Thus, our work examining LLR in young animals and in a model targeted at severe lung injury, may succeed where liquid ventilation for adult ARDS did not.

Unlike liquid ventilation, LLR does not necessarily require the gas exchange properties of perfluorocarbons. In most cases of ECLS, oxygenation and carbon dioxide removal can be fully performed by the extracorporeal oxygenator, pulmonary gas exchange. Thus, LLR opens the possibility of using liquids that do not perform efficient gas transfer. However, there are several benefits of using PFOB over other fluids, such as saline. First, its high density (∼2 g/cm^3^) and low viscosity (∼3 centistokes), allow it to flow to dependent portions of the lung, recruiting alveoli and displacing mucus, pathogens, and inflammatory cells (16). Second, PFOB has a high capacity to dissolve gas, which allows it to absorb any trapped alveolar air, preventing potentially harmful air pockets. Third, PFOB evaporates at 1-2 ml/kg/hr, offering a seamless transition back to conventional gas ventilation without surfactant washout. Finally, in the event of an ECLS device failure, PFOB allows the lungs to be used for gas exchange. Although PFOB has several benefits, exploration of other potential liquids (or mixtures with PFOB) with antimicrobial, immunomodulatory, or other beneficial biological effects may yield improved results.

A key finding of this study was the large improvement in alveolar opening in the most dependent regions of the lung. In zone 6 (posterior region of the lower lobe), alveolar airspace was more than double in the LLR group compared to the gas group. This dramatic treatment effect is likely a combination of 1) the tendency of the dependent zones to collapse in ARDS and 2) the dense nature of PFOB, allowing it to flow to the most dependent regions. Of note, our study utilized a relatively small dose of PFOB (5 mL/kg). Prior studies of PFOB in liquid ventilation have utilized doses between 2.5 mL/kg and 30 mL/kg (29, 33). We chose a low dose due to concerns from clinical trials that pulmonary overdistension with excess PFOB affect hemodynamics. Our study suggests that 5 mL/kg may be too little to achieve alveolar opening throughout the lung. Follow-up studies to identify the optimal PFOB dosing for LLR are warranted.

We had mixed results with regards to the effect of LLR on inflammation. We did not find significant differences in plasma cytokine levels. This is somewhat expected, as the relatively short time course (4 hours) likely meant that plasma cytokine levels were primarily driven by the acute lung injury and initiation of ECLS, rather than the differential impact of liquid versus gas lung rest on biotrauma. Additionally, our study was powered for differences in C_stat_, as opposed to cytokine levels, which may have led to Type II error. Furthermore, one outlier pig in the gas group was excluded, which biased our results toward the null hypothesis (i.e. no difference in cytokine levels), though sensitivity analyses including the pig did not change our conclusions. Despite no significant differences in most cytokine levels, an interesting finding was the difference in IL-10 concentration in the lung tissue, even after just 4 hours. IL-10 is a key anti-inflammatory cytokine and has been suggested as a potential therapy for ARDS (35, 36).

However, despite its immune regulatory role, multiple studies have identified IL-10 as an early biomarker portending a poor prognosis in ARDS (37-40). Liu et al. measured IL-10 levels in ARDS patients receiving ECLS and found that IL-10 levels at the time of cannulation and during the first 6 hours of ECLS support were significantly higher in patients that died compared to survivors (39). In our study, the lower levels of IL-10 in the LLR-treated animals could be due to a direct anti-inflammatory effect of PFOB or reduced ongoing lung injury.

Our study has several limitations. First, while the oleic acid lung injury model has been utilized for decades by multiple research groups, its mechanism of injury mimics a fat embolism, which is a rare trigger of lung injury in humans. Second, our injury model only achieved moderate ARDS, with an average P/F ratio of 182. Although additional doses of oleic acid could achieve a more severe injury, doses higher than 0.12 mL/kg caused severe hemodynamic compromise, which often led to the premature death of the pig. Other injury models such as saline lavage or direct acid injury were considered (41), though these are limited by a high variability in the degree of injury. Additionally, ARDS is a dynamic disease that evolves over time through proteinaceous fluid infiltration into the alveoli, surfactant inactivation, and ongoing secondary injury from mechanical ventilation and immune activation. While our model achieves a significant acute lung injury, a longer time course is necessary to reproduce severe ARDS. The 4-hour time course also limits our ability to draw conclusions about lung recovery during LLR. Our findings of improved compliance, dependent zone alveolar opening, and reduced heterogeneity suggests that LLR may reduce ongoing atelectrauma and enable optimal lung rest. However, whether this translates to faster lung recovery over days to weeks remains an unanswered question for future work.

## 5. Conclusion

ECLS is a life-saving therapy for severe ARDS that can allow for lung rest, but the optimal lung rest strategy is unknown. In this study, we found that in comparison to standard gas ventilation for lung rest, LLR can improve compliance and increase alveolar recruitment, especially in the most dependent areas of the lung. LLR offers a potential therapy that may improve lung recovery in patients with severe ARDS requiring ECLS.

