## Supplemental Material for "Liquid Lung Rest During Extracorporeal Life Support in a Porcine Model of Acute Lung Injury"

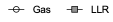

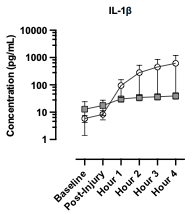

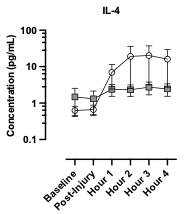

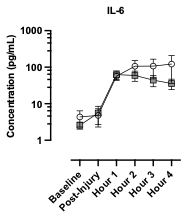

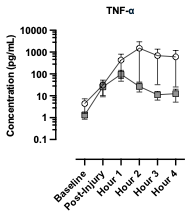

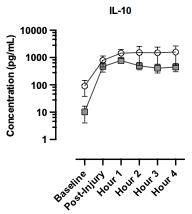

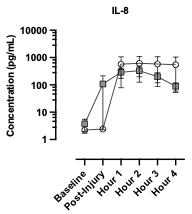

*

*

*

*

*

*

*

*

*

*

*

*

*

*

*

*

*

*

*

*

*

*

*

*

*

*

*

*

*

*

*

*

*

*

*

*

*

*

*

*

*

*

*

*

^†^

**Figure S1. Plasma cytokine levels (all pigs included)**

Plasma samples were drawn at baseline, after lung injury, and hourly while on ECLS. These data incorporate all pigs, including an outlier pig from the gas ventilation arm. Y-axis is presented on a logarithmic scale. Error bars denote standard error.

p<0.05, ⁎: versus baseline value; †: versus gas ventilation

**Concentration (pg/mL)**
